# InterPepScore: A Deep Learning Score for Improving the FlexPepDock Refinement Protocol

**DOI:** 10.1101/2021.12.09.471890

**Authors:** Isak Johansson-Åkhe, Björn Wallner

## Abstract

**Motivation:** Interactions between peptide fragments and protein receptors are vital to cell function yet difficult to experimentally determine the structural details of. As such, many computational methods have been developed to aid in peptide-protein docking or structure prediction. One such method is Rosetta FlexPepDock which consistently refines coarse peptide-protein models into sub-Ångström precision using Monte-Carlo simulations and statistical potentials. Deep learning has recently seen increased use in protein structure prediction, with graph neural network seeing use in protein model quality assessment.

**Results:** Here, we introduce a graph neural network, InterPepScore, as an additional scoring term to complement and improve the Rosetta FlexPepDock refinement protocol. InterPepScore is trained on simulation trajectories from FlexPepDock refinement starting from thousands of peptide-protein complexes generated by a wide variety of docking schemes. The addition of InterPepScore into the refinement protocol consistently improves the quality of models created, and on an independent benchmark on 109 peptide-protein complexes its inclusion results in an increase in the number of complexes for which the top-scoring model had a DockQ-score of 0.49 (Medium quality) or better from 14.8% to 26.1%.

**Availability:** InterPepScore is available online at *http://wallnerlab.org/InterPepScore*.

## 1 Introduction

Interactions between a protein receptor and a smaller flexible peptide fragment make up 15-40% of all protein-protein interactions (Petsalaki and Russell, 2008), and are involved in vital cell functions such as cell life-cycle regulation (Midic *et al*., 2009). The flexibility of the peptide and often transient nature of the interaction (Tu *et al*., 2015) makes them difficult to study experimentally. Thus, computational proteinpeptide docking methods are needed to understand the molecular mechanisms and implications of the interactions (Petsalaki and Russell, 2008; Helander *et al*., 2015; Wei *et al*., 2019).

The approaches to solving the protein-peptide docking problem ranges from advanced searches for structural templates of interaction such as with InterPep2 (Johansson-Åkhe *et al*., 2020) or SPOT-peptide (Litfin *et al*., 2019), to exhaustive sampling of docked conformations as in PIPER-FlexPepDock(Alam *et al*., 2017) or pepATTRACT (Schindler *et al*., 2015), and end-to-end machine-learning generated models of interaction as with the input adapted AlphaFold2 protocols (Tsaban *et al*., 2021). Common to most of the docking approaches is the need for a final step of all-atom refinement to optimize the final structures. Several of the methods above use the Rosetta Flex-PepDock refinement protocol (Raveh *et al*., 2010) for this final step.

Here, we introduce the InterPepScore scoring term for use with the Rosetta FlexPepDock refinement protocol. InterPepScore is a graph neural network trained to predict the resulting DockQ score (Basu and Wallner, 2016) during FlexPepDock Monte Carlo sampling. When added as an additional scoring term, it consistently improves the performance by generating more high quality models.

## 2 InterPepScore – Development and Performance

### 2.1 InterPepScore Training

The InterPepScore scoring term was trained and validated on Rosetta FlexPepDock simulation trajectories, including both accepted and rejected structures during the Monte Carlo sampling. Trajectories were generated from initial coarse models of 4,447 peptide-protein complexes, by perturbing the native peptide (Raveh *et al*., 2010), template-based docking (Johansson-Åkhe *et al*., 2020), or rigid-body docking (Alam *et al*., 2017) (see Supplementary Methods for details). Online training of InterPepScore was performed by iteratively generating new FlexPepDock trajectories using the FlexPepDock protocol including InterPepScore as a scoring term and adding these new trajectories to the training data. This was done whenever training converged to augment the training set and ensure that the final model was trained on data similar to what it would generate. InterPepScore was implemented using pyRosetta (Chaudhury *et al*., 2010).

### 2.2 Graph Network Architecture

InterPepScore is a global score implemented as a graph network with protein residues as nodes and edges between residues within 10 Å. The node features are BLOSUM62 columns representing the amino acid residues and one-hot identifiers if the residues belong to the protein or peptide chain. BLOSUM62 was used instead of sequence embedding as only marginal improvements can be found for modest sized data set, and only after many epochs of training (ElAbd *et al*., 2020). Multiple sequence alignments were not used to ensure InterPepScore can be used as any other Rosetta scoring term without additional pre-calculation of features. The edge features denote only if the residues are covalently bound or not and if the edge is a self-edge.

Three layers of graph network forward passes - updating all node, edge, and global features with ELU activation as implemented through DeepMind’s Graph Nets package (Battaglia *et al*., 2018) followed by a final convolution of the global features make up the architecture of the network. During training, dropout and batch normalization were used. More details on the network architecture can be found in the Supplementary Information.

### 2.3 InterPepScore improves refinement

The performance of the FlexPepDock refinement protocol when including the InterPepScore scoring term was analyzed on a test set of 109 non-redundant peptide-protein complexes that shared no CATH superfamily annotation with any complexes from the training or validation sets. For each complex, starting points were generated the same way as for the training data For the test and validation sets, each method for generating starting points contributed with four starting models, in total 12 starting points per complex. The FlexPepDock refinement protocol was run 200 times with and without InterPepScore for each starting point. In all cases, the receptor was relaxed and the exposed side-chains perturbed prior to docking to simulate an unbound-to-bound docking scenario.

By including the InterPepScore both in the sampling and selection stage of the refinement the DockQ score is improved (Figure 1), e.g. with the original FlexPepDock protocol 14.8% of the targets had a DockQ indicating Medium quality model or better, this is improved to 26.1% of the targets using InterPepScore. The original FlexPepDock refinement protocol already consistently improves the quality of all decoys except those with high DockQ scores. But with InterPepScore the improvement is larger and maintained even for decoys with already excellent starting DockQ (Figure S1). InterPepScore not only aids in final selection, but also leads to FlexPepDock producing higher quality models overall (Figure S1 and S2).

**Fig. 1:**
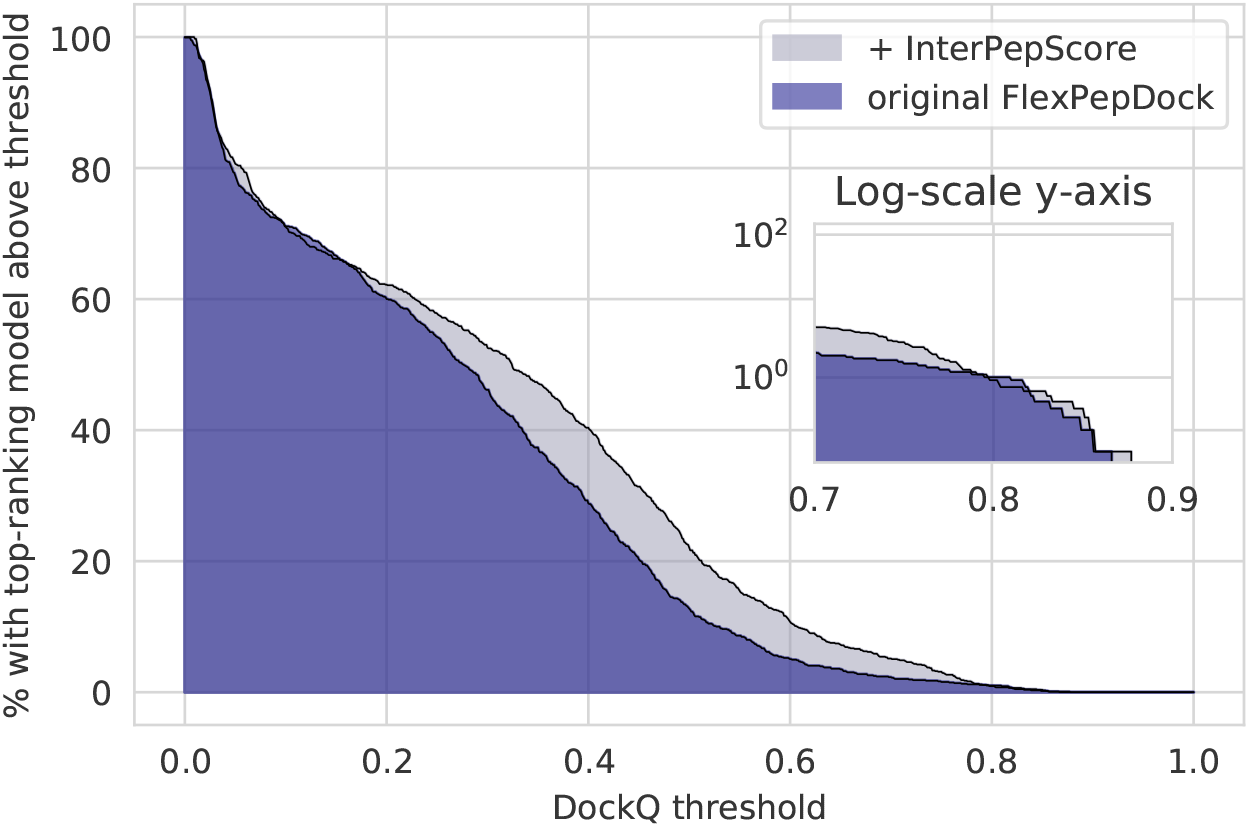
For the 109 peptide-protein complexes investigated, the addition of InterPepScore to the FlexPepDock refinement protocol both during folding and final decoy selection consistently improves the quality of the final selected decoys as measured by DockQ score (Basu and Wallner, 2016). With InterPepScore, the top-scoring decoy achieves a DockQ-score of at least Medium quality (0.49) in 26.1% of all cases, as opposed to only 14.8% without.

### 2.4 InterPepScore refines AlphaFold models

To demonstrate general usefulness of InterPepScore in refinement, we generated starting models for the test set using AlphaFold with a 30 glycine linker (Tsaban *et al*., 2021). Even though the improvements in absolute terms are small and most refinements (85/109) do not change the DockQ score significantly, InterPepScore still improve 19/109 targets significantly, while only 5/109 targets get significantly worse (Figure S3).

## 3 Conclusion

InterPepScore is easy to use with pyRosetta and contributes significant improvements towards the FlexPepDock refinement protocol. It is our hope that many of the state-of-the-art peptide-protein docking pipelines which make use of this refinement protocol will also make use of this additional scoring term to improve their refinement steps.

## Acknowledgments

This work was supported by SeRC, VR 2020-03352, CTS 20:453. Computations provided by SNIC, KAW, and LiU through NSC.

## Notes

### Competing Interest Statement

The authors have declared no competing interest.

