## Supplementary for "InterPepScore: A Deep Learning Score for Improving the FlexPepDock Refinement Protocol"

### InterPepScore Supplementary Information

Isak Johansson Åkhe    Björn Wallner

December 9, 2021

### 1 Method

#### 1.1 Data set

To train and analyze the method, peptide-protein complexes from the PDB (Berman *et al.*, 2000) fetched at 20/12 2018 were redundancy reduced by clustering at 30% sequence identity down to 687 complexes. In this case, a peptide-protein complex was considered any protein-protein complex where one ”receptor” was at least 50 residues and shared at least 200 Å$^2$ of contact surface with a ”peptide” chain of less than or equal to 25 residues.

A set of 109 complexes were selected as a test set and a separate set of 95 complexes selected as validation set. The sets were generated randomly, but if any receptor of any set shared a CATH superfamily annotation with any other set (including the remaining 480 complexes), the randomization was repeated until the sets shared no such connections. In the case that a receptor had no CATH annotation, it was awarded the same annotation as the chain in the full non-redundancy-reduced set that it matched to with the greatest TM-score when aligned with TM-align (Zhang and Skolnick, 2005).

The training set was constructed by including every complex from the initial non-redundancy reduced set which did not share a CATH superfamily annotation with neither the test set nor the validation set, resulting in 4,447 peptide-protein complexes used for training.

#### 1.2 Generating initial starting structures

The initial coarse models used as starting points for the FlexPepDock trajectories were generated using three different schemes: (i) by perturbation of the native peptide similarly to the original FlexPepDock refinement paper (Raveh *et al.*, 2010), (ii) template-based docking similarly to InterPep2 (Johansson-Åkhe *et al.*, 2020), and (iii) rigid-body-docking with several sampled peptide conformations similarly to PIPER-FlexPepDock (Alam *et al.*, 2017). See below for details.

For the test and validation sets, each method for generating starting points contributed with four starting models, in total 12 starting points per complex.

For the training set, a total of 440,000 starting points were generated evenly distributed between the CATH superfamilies of the training set. Half of all starting points were generated by rigid-body-docking, and the rest were evenly distributed between perturbation from the native fold and template-based docking, respectively.

To ensure having enough starting points close to the correct structure to enable refinement, at least half of all starting points from each generation method needed to have an LRMSD $< 5.5$ Å or any contiguous stretch of at least half the peptide (or 5 residues, whichever is larger) with an LRMSD $< 4.0$ Å. The first cutoff was chosen as it is the limit for salvageable decoys from the FlexPepDock (Raveh *et al.*, 2010). For longer peptides, sometimes more of the peptide is modeled than what is actually in contact with the receptor, which is why the second cutoff was also allowed. The forced selection was obained by continually discarding generated positions and generating new ones until the thresholds were satisfied.

Details for how starting points were generated using the different methods can be found below:

(i) **Perturbation of the Native Position:** PyRosetta was used to apply random shear, small, rigid-transformation, and rigid-rotation movements to the native peptide. In this starting point generation scheme, the size of the changes applied were modified after each starting point generated in accordance with if that point became salvageable or unsalvageable (increased movement sizes if the point became salvageable and vice versa).

(ii) **Template-based Docking:** The receptor was aligned by TM-align to every protein-protein complex in the PDB (Zhang and Skolnick, 2005). The same rigid configurations for the peptide as used for the rigid-body docking were structurally aligned to the other side of the interface for the best scorers by use of InterComp (Mirabello and Wallner, 2018). This scheme of docking is similar to that of InterPep2, but without the random forest evaluation (Johansson-Åkhe *et al.*, 2020).

(iii) **Rigid-Body Docking:** Rigid configurations of the peptide was generated as in the PIPER-FlexPepDock paper and docked with PIPER on the receptor surface (Alam *et al.*, 2017). The best-scoring decoys by PIPER score were selected.

#### 1.3 Network Architecture

An overview of the graph network architecture can be found in Figure S1. The three Graph Network blocks are the same as the suggested graph network architecture from (Battaglia *et al.*, 2018), with the learning component (blue blocks of the figure) composed of the following, in feed-forward order:

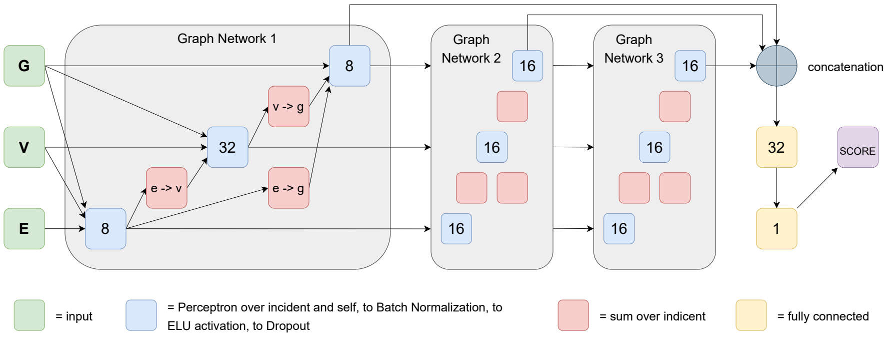


Figure S1: The neural network architecture of InterPepScore. E, V, and G represents edge features, node (vertex) features, and global features respectively. The blue boxes represent the core learning elements of the network. The numbers inside boxes indicate the length of the feature vector they output. Note that in a graph network, the V row always outputs one feature vector *per node*.

1. **Concatenation** of the feature vectors of all incident elements and the active elements' own feature vectors. In cases where the number of incident elements are unknown, the blue boxes are always preceded by a red box, which indicates a summation of the feature vectors of the incident elements. The global feature vector is incident to all other elements.
2. **Perceptron**: The basic learning component is a simple linear perceptron.
3. **Batch Normalization**
4. **ELU activation**
5. **Dropout:** During training, a 40% dropout rate of features was employed.

#### 2 Extended Analysis

Figure S2 shows that the FlexPepDock refinement protocol (Raveh *et al.*, 2010) successfully produces and selects decoys with better DockQ-scores (Basu and Wallner, 2016) than their starting points for all cases except those where the starting point is already modeled with sub-Ångström precision. Additionally, in the same figure it is evident that the addition of InterPepScore further improves upon this, and does not disturb the peptide when already modeled with great precision.

FlexPepDock with or without InterPepScore frequently generates decoys with worse DockQ-scores than their starting positions, when all generated decoys are considered, and not only the top decoys selected using the best scores, Figure S3. This is expected as FlexPepDock is a Monte-Carlo based approach

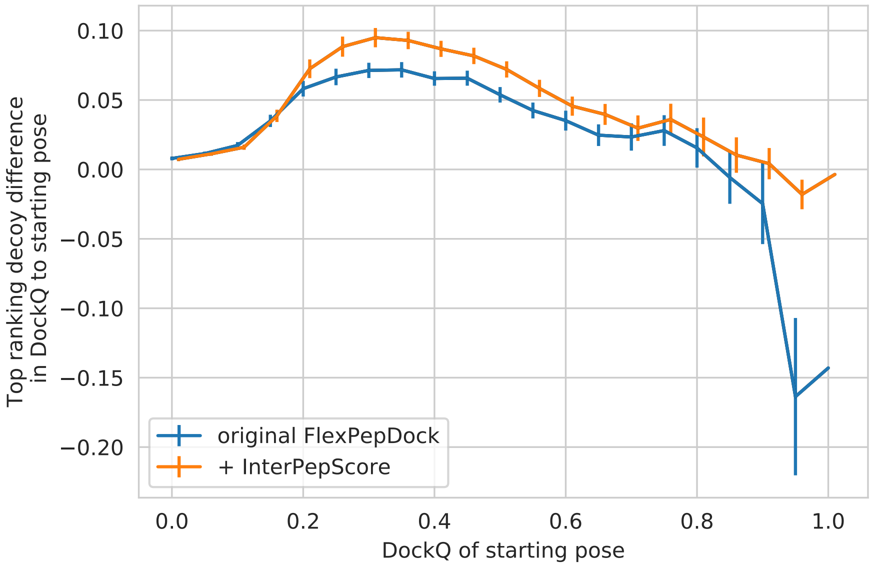


Figure S2: For each starting position for each of the 109 peptide-protein complexes investigated, the addition of InterPepScore to the FlexPepDock refinement protocol both during folding and final decoy selection consistently improves the quality of the final selected decoys as measured by DockQ score.

which often end up in local minima or unfavorable positions, requiring the protocol to be run many times and relying on the scoring function to select structures similar to a native structure. Also visible in the same figure is the fact that InterPepScore generates a much higher contrast between poor- and high-quality final decoys (the coloring). The coloring of the graph is linear between lowest and highest value, but outliers were excluded in calculation of extreme values by iteratively dropping data points further than 3 standard deviations from the mean.

### 3 FlexPepDock refinement with InterPepScore of AlphaFold2 models

In the main paper, using FlexPepDock refinement with InterPepScore is tested on decoys generated by template-based docking, rigid-body docking, and perturbation of the native peptide.

To test the applicability of FlexPepDock refinement with InterPepScore on models generated by the state-of-the-art simultaneous folding and docking protocol utilizing AlphaFold2 proposed by (Tsaban *et al.*, 2021), this implementation was used to generate starting points for the same 109 complexes considered

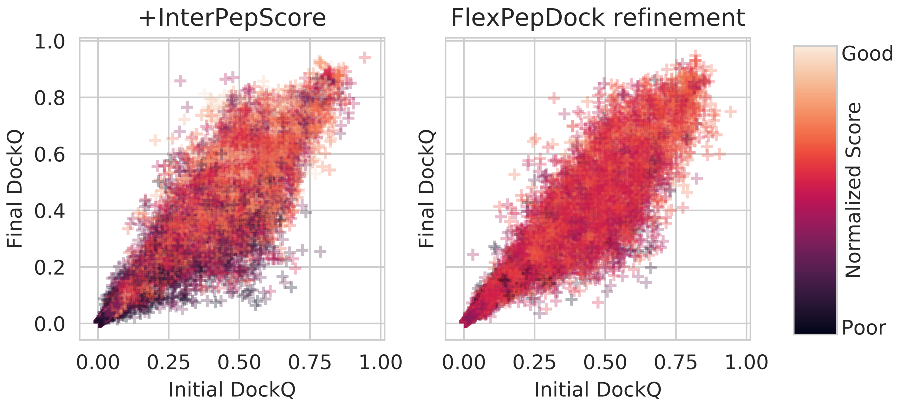


Figure S3: For every single run of FlexPepDock for every starting position, the DockQ-score of the decoy produced compared to its starting position. Graphs are colored by the metric used for evaluating the final complex; InterPepScore when it was included as a scoring term and the reweighted-score scoreterm traditionally used to rank decoys with FlexPepDock refinement. The reweighted-score scoring term is normalized by protein size and the outliers are pruned from the score span specified in the method description.

in the main text. In summary, the AlphaFold2-based docking involves using a polyglycine linker between the receptor and peptide to submit the complex to the AlphaFold2 inference step as one single protein chain.

As can be seen in Figure S4, using FlexPepDock refinement with InterPepScore improves the quality of these models much like for the coarse models investigated in the main text (compare with Figure S2).

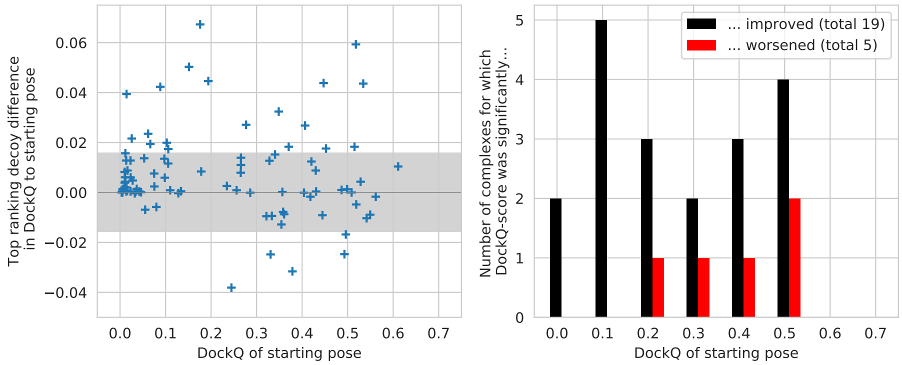


Figure S4: The capacity for FlexPepDock refinement including the InterPep-Score score term to improve the quality of models created by using AlphaFold2 with a polyglycine linker as proposed in Tsaban *et al.* (2021). Points outside the shade area are significantly (P<0.05) changed, positive differences are improved relative to the starting pose.
